# A Machine Learning and Bioinformatic Analysis Reveals an Associated between Cell Surface Receptor Transcript Levels with Drug Response of Breast Cancer Cells and the Drug Off-Target Effects

**DOI:** 10.1101/2022.08.31.506005

**Authors:** Musalula Sinkala, Krupa Naran, Dharanidharan Ramamurthy, Neelakshi Mungra, Darren Martin, Stefan Barth

**Author notes:** **Corresponding author’s e-mail addresses** Bioinformatics, Precision Medicine.

## Abstract

Breast cancer is characterised by varied responses to different anticancer therapies, which may provoke several different off-target effects. We hypothesise that for drugs that target cell surface receptors (CSRs), the different responses of tumours and the adverse events produced by these drugs may be attributed to variations in the transcriptional landscapes of CSRs in both breast tumours and healthy tissues. Here, we use data from various sources to compare the CSR transcriptional landscapes of breast tumours and a range of different non-diseased human tissues. We demonstrate an association between the responses to drug perturbation of breast cancer cell lines and the transcription levels of their targeted CSRs. Furthermore, we reveal important differences in the CSR transcriptional landscapes of primary breast tumour subtypes and the CSR transcriptional landscapes of breast cancer cell lines, which will likely impact the accuracy of drug response predictions. Finally, applying clinical trial data, we expose a link between the expression levels of CSR genes in healthy tissues and adverse reactions of patients to anticancer drugs. Altogether, this approach allows for the isolation of the most suitable CSR target(s) among the expressed transcripts, solely based on the measured dose-responses of cell lines to small molecules, the CSR transcriptional landscape in health patient tissues, and reported adverse responses of patients to drugs targeting CSRs.

## Introduction

Among the most notable differences between cancer cells and healthy untransformed cells is the differential overexpression of particular cell surface receptor (CSR) proteins on cancerous cells. CSR proteins traverse the plasma membrane to provide sensory links between the signalling pathways of the cytosol and the extracellular environment. Besides being exposed on the external surfaces of cells, alterations in the signalling pathways within which many CSRs function, are directly involved in oncogenesis. CSRs are, therefore, often effective targets for anticancer drugs and antibody-based anticancer therapies [1–4]. Alterations in the expression of CSRs are widespread during oncogenesis and can involve gene mutations, gene copy number changes and/or transcriptional changes [5–7].

CSRs currently targeted to treat tumours of the breast and other tissues were initially identified as useful targets based on either their differential overexpression in tumours compared to their adjacent healthy tissues or their mutational profiles in cancer cells [8–10]. However, this approach to identifying credible targets has generally given little consideration to the expression levels of targeted CSRs in the tissues of body organs that are not directly associated with the primary tumours under consideration.

Numerous clinical trials that aim to evaluate the efficacy of CSR-targeted treatments ultimately fail due to the dose-limiting toxicity and unexpected side effects [11–13]. Many of these undesirable side effects likely result from interactions between drugs or therapeutic antibodies and CSRs in healthy tissues [14,15]. If so, treatment success rates could be improved, and off-target toxicity could be reduced by targeting CSRs that are both overexpressed in cancer cells and under-expressed in all healthy tissues.

Here, we attempt to identify such CSRs in the context of finding targets for breast cancer treatment. Specifically, we provide a platform for hypothesis generation and a framework for selecting which CSRs should be targeted to minimise the probability of undesirable treatment side effects. We mine and integrate data from multiple public resources and apply statistical methods, machine learning and predictive modelling to investigate the probable off-target toxic effects of targeting a variety of different CRSs; including many that are currently targeted and for which actual toxicity effects have been measured. We make several explicit assumptions regarding the relationships between drug action in the context of drug responses and off-target toxicity, and the transcriptional landscapes of the breast cancer cells and those of healthy body tissues and evaluate these concepts using data from thousands of high-quality laboratory measurements.

Here, we use a bioinformatics approach to investigate, in breast cancer cell lines, the relationship between the transcription levels of the CSRs and the cell line’s responses to drugs that targets specific CSRs. Furthermore, we evaluate the association between the adverse (or off-target) effects of CSR-targeted drugs that are used to treat breast tumours and the transcriptional landscapes of the targeted CSRs in various healthy body tissues. Overall, our computational approach reveals the link between drug action and the expression of CSRs in breast tumours, an insight that is possibly valid across many cancers of other tissues and provides a framework for improving the criteria by which drug targets are selected.

## Results

### The transcriptional landscape of CSRs across breast tumours and healthy tissues

To define patterns of CSR transcription across various normal tissues, we retrieved and collated mRNA transcription data of 101 major body organs and tissues from the GTEx project [16–18]. Only the mRNA transcription signatures of 1,140 genes that encode CSR proteins were returned, on which we applied unsupervised hierarchical clustering to investigate variations in the expression of CSRs across different healthy organs and tissues (Figure 1A). We then used similarly processed mRNA transcription data from breast cancer samples obtained from the cancer genome atlas (TCGA) [19] (1,091 samples) and compared these with CSR Mrna transcription data from healthy tissues obtained from the GTEx project (9,658 samples, including 218 samples from healthy breast tissue; Figure 1B and 1C).

**Figure 1:**
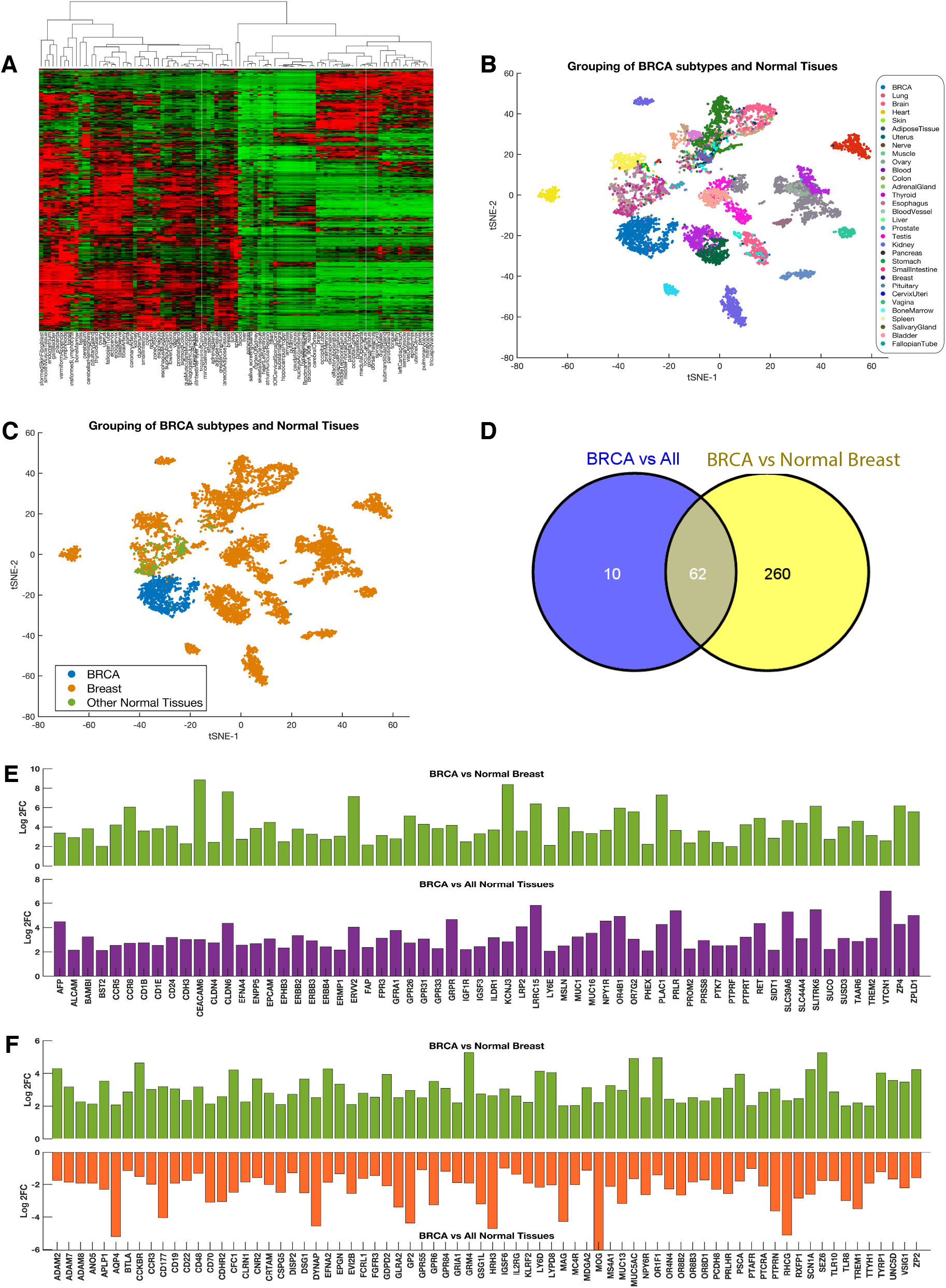
(A) Clustered heatmap of 101 normal human tissues using the mRNA transcription data of CSR. The heatmap was produced using unsupervised hierarchical clustering with the Euclidean distance metric, row standardisation and complete linkage. (B) Clustering of BRCA samples (blue points) and normal tissue samples based on mRNA transcript levels of CSRs. Points are coloured according to the tissues from which the measured CSR transcripts were obtained. (C) Clustering of 1,320 BRCA samples (blue points), 13,390 normal breast tissue samples (green points) and 34,000 other normal tissue samples (orange points) based on mRNA transcript levels of CSRs. For both figure panels (B and C), t-SNE was used to visualise the sample clustering using the exact algorithm and standardised Euclidean distance metric. (D) The number of CSR transcripts upregulated between breast tumours vs all other normal tissues and breast tumours vs normal breast. Sixty-two transcripts are commonly upregulated between the two sets of comparisons. Refer to supplementary file 1 and 2 for the complete results of the differentially expressed CSR transcripts. (E) Bar graph showing CSR transcripts that are commonly upregulated for the two comparisons: 1) between breast tumours vs normal breast and 2) between breast tumours vs all other healthy tissues. (F) CSR transcripts that show a reverse signature. These are upregulated in breast tumours compared to the healthy breast, but downregulated breast tumours compared all other normal tissues.

We identified 634 CSR transcripts that were differentially expressed (4-fold-change > 2 or < −2, and adjusted p-value < 0.05) between the breast tumours and healthy breast tissue, and 581 CSR transcripts that were differentially expressed between breast tumours and healthy body tissues in general. Here, we found 322 CSR transcripts that were more highly expressed in breast tumours than in healthy breast tissue (Figure 1D). Among the most significantly up-regulated transcripts were CEACAM6 (log2FC = 8.8), KCNJ (8.4) and CLDN6 (7.6; see Supplementary File 1). Furthermore, we found 72 CSR transcripts that were significantly upregulated in breast tumours compared to non-breast healthy body tissues (Figure 1D). Among these were VTCN1 (log2FC = 7.0), LRRC (5.8) and SLITRK6 (5.5). Here, we found that only 62 transcripts were shared between those that were: (1) upregulated in breast tumours relative to the healthy breast tissue and (2) upregulated in breast tumours relative to healthy non-breast tissues (Figure 1D). Included among these are transcripts for ERBB2, ERBB3, EPCAM, and IGFR (Figure 1E).

Among the 511 CSR genes that were upregulated in breast tumours compared to healthy breast tissues, we found 72 significantly downregulated in breast tumours compared to healthy non-breast tissues. Among these 72 transcripts are well-known drug targets used in treating breast cancer, including FGFR3, CD48 and CCR3 (Figure 1F). These findings emphasise that even when targeted CSRs are more highly expressed in breast tumours than in healthy breast tissues, it does not necessarily entail that they will also be more highly expressed in breast tumours than in other healthy tissues.

Natural inter-tissue variations in the expression of CSR proteins, such as those highlighted above, are expected to complicate the translation of CSR-binding anticancer molecules into useful therapeutics. Specifically, the presence of large numbers of targeted CSRs on the cells of healthy tissues is likely to be the primary cause of the dose-limiting toxic effects that are commonly associated with CSR-targeted anticancer drugs [14,15].

### Identification of Ideal Targets

Accordingly, we hypothesised that such off-target toxic effects could be ameliorated by targeting CSRs expressed at higher amounts on cancer cells than on all healthy breast and non-breast tissue types.

Here, we considered if a threshold level of mRNA expression could be determined to aid in ranking CSRs. Overall, we determined the difference in the mean expression levels between breast tumours and healthy breast tissues. Therefore, to designate a given target, we used an approach taking into account the expression of these targets across all healthy tissues such that essential tissues (brain, heart, lung, liver and/or kidney) should be avoided while toxicities in non-essential tissues should be minimized (see methods sections). Briefly, we looked for CSR whose mRNA transcriptional levels were significantly upregulated in breast tumours compared to any other healthy tissues. By comparing CSR transcript levels in breast tumours to those in all healthy tissue types, we identified 26 CSRs that, by being substantially more highly expressed in breast tumours than in any healthy tissue type, could potentially be targeted with minimal off-target cytotoxic effects (Figure 2, also see Supplementary File 1 and 2).

**Figure 2:**
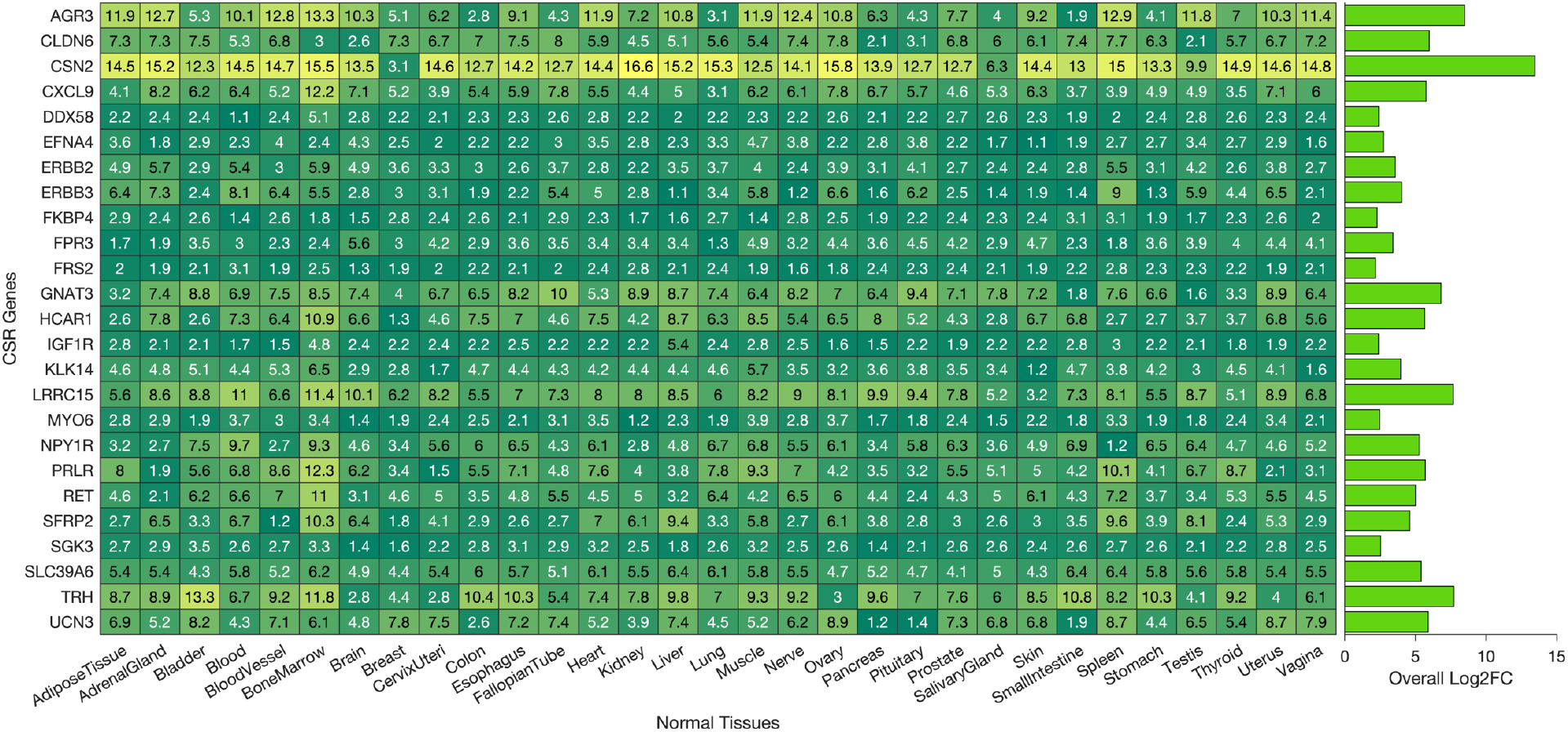
The heatmap of the log-2 fold-change values of the CSR transcripts that are consistently upregulated in our comparison of breast tumours against every other healthy tissue.

### PAM50 breast cancer subtypes exhibit distinct CSR transcriptional patterns

Breast tumours are classified into five molecular subtypes based on a 50-gene signature called PAM50 [20,21]. These PAM50 subtypes are Luminal A, Luminal B, Normal-like, Basal-like and HER-2 positive. Here, by comparing the CSR transcript levels between tumours annotated as belonging to the different PAM50 subtypes by the TCGA, we found substantial differences in the transcript levels of various CSR genes between the subtypes (Figure 3A, Supplementary File 3). We found the highest number (323) of differentially expressed transcripts between Basal-like and Luminal A breast tumours and the fewest (32) between Luminal A and Luminal B breast tumours. Overall, our results show that Basal-like tumours have significantly different transcription profiles from those of Luminal A and Luminal B tumours, of which the transcriptional signatures of Luminal A and B tumours are relatively more similar.

**Figure 3:**
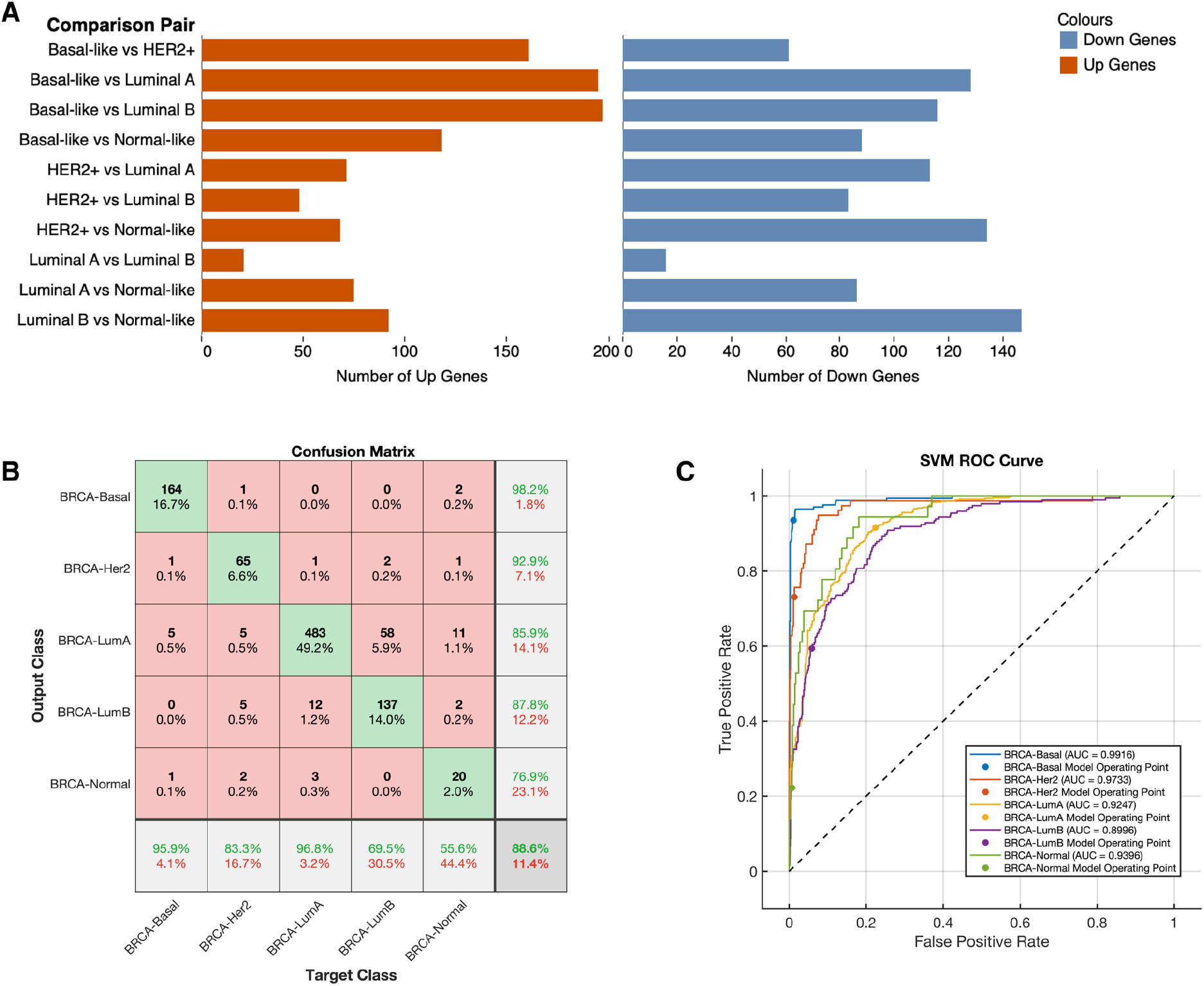
(A) Bar graphs showing the number of differentially expressed CSR transcripts between each pairwise comparison of the PAM50 breast cancer subtype (x-axis). The two bar graphs are plotted for the upregulated transcripts between each comparison (left) and downregulated transcripts between each comparison (right; also see Supplementary File 3). (B) A plot of the confusion matrix. The diagonal cells correspond to correctly classified observations (green cells). The off-diagonal cells correspond to incorrectly classified observations (red cells). Both the number of observations and the percentage of the total number of observations are shown in each cell. The column to the far right shows the precision (or positive predictive value rate) in green text and the false discover rate shown in red text. The row at the bottom of the plot shows the recall (or true-positive rate) in green text and the false-negative rate in red text. The cell in the bottom right of the plot shows the overall accuracy. (C) shows the ROC-AUC (Receiver Operated Characteristic-Area Under the Curve) for the PAM50 breast cancer classes. The ROC curves show the true-positive rate versus the false-positive rate for predictions of the classes made by the trained classifier. The coloured markers on the plots show the values of the false positive rate and the true-positive rate of the trained classifier toward predicting the PAM50 subtype of breast cancer using CSRs.

Since PAM50 subtyping of the breast tumours is critical in the treatment and prognosis of breast cancer, we sought to evaluate whether variations in CSR mRNA transcript levels alone could be utilised to classify breast cancer tumours accurately. Therefore, we used the PAM50 annotations provided within the TCGA database for each sample and the CSR transcription data to train a supervised machine learning model (see methods sections) to classify tumours.

Here, by applying an ensemble boosted decision tree model, we found that we could accurately predict (average area under the curve of 93% and classification accuracy of 89%) the PAM50 subtype of breast tumour samples based exclusively on 1,140 CSR transcript levels (Figure 3B and 3C). Furthermore, our model had positive predictive values of 85.9% for Luminal A, 87.8% for Luminal B, 76.9% for Normal-like, 98.2% for Basal-like and 92.9% for HER-2 positive breast cancer (Figure 3). Overall, these results indicated that HER-2 and Basal-like breast tumours have the most distinctive CSR transcription profile, whereas Luminal A and Luminal B have transcription profiles that are the most difficult to differentiate between.

### Drug responses are associated with the transcriptional levels of targeted CSRs

We sought to evaluate whether the response profiles of breast cancer cell lines to CSR-targeted drugs differed in association with the PAM50 subtype classification of the cell lines. Using drug response data from the GDSC [22] for Luminal A, Luminal B, Basal-like and HER2 breast cancer cell lines, we made a pairwise comparison of the mean drug responses between cell lines of each breast cancer subtype to thirteen CSR targeting drugs (Figure S1). After correcting for multiple comparisons, we found that, except for Basal-like versus Luminal A breast tumours (adjusted p-value = 0.0079), the drug-response profiles of the different subtypes of cancer cell lines were not significantly different (see Supplementary File 4).

We hypothesised that the transcriptional profiles of CSRs of the breast cancer cell lines that are represented within the GDSC database might differ from the transcriptional profiles of primary tumours (such as those represented within the TCGA database). If so, it would be expected that, for a particular CSR-targeting drug, drug responses of the cell lines would vary based primarily on differences between the expression levels of the targeted CSR.

To test our hypothesis, we focused our analysis on CSR transcripts that are consistently differentially expressed between cell lines classified as belonging to different PAM50 subtypes.

We found no CSR transcripts at significantly different levels (adjust p-values less than 0.05 and 2-fold change) higher than one or less than minus one) when comparing the Basal-like and HER2+ subtypes or the HER2+ and Luminal B subtypes. In addition, we found only one CSR transcript with significantly different levels in the Basal-like and Luminal B subtypes of breast cancer cell lines. Therefore, relative to primary tumours belonging to the different PAM50 subtypes – which display substantial differences in CSR expression (Figure 4 and Supplementary File 4) – it appears that breast cancer cell lines display far more uniform CSR expression patterns across the subtypes. This may explain, at least in part, the observed uniformity in responses to CSR-targeted drugs across breast cancer cell lines belonging to different PAM50 subtypes.

**Figure 4:**
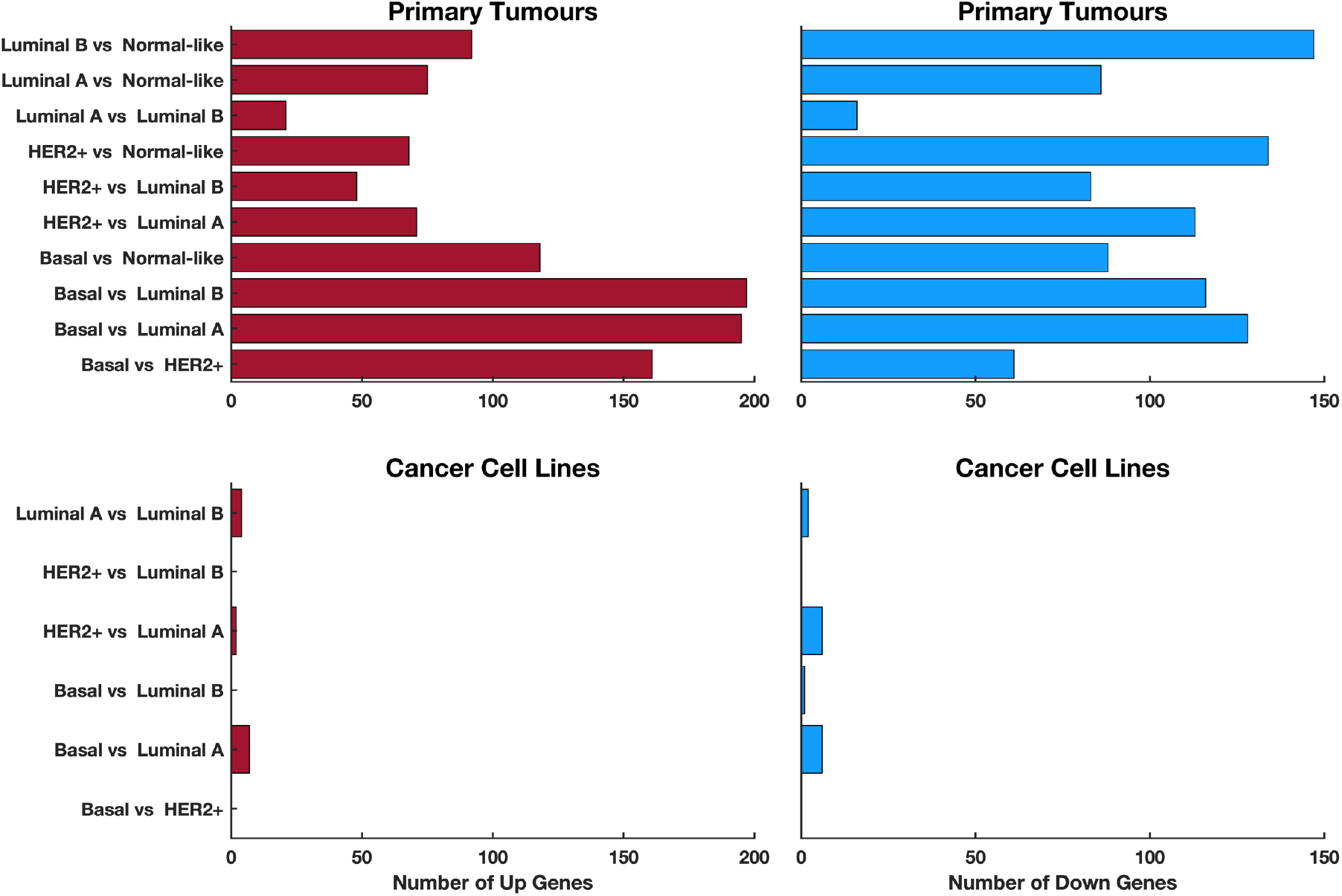
Comparison between the number of differentially expressed CSR transcripts between each pair of PAM50 subtypes of breast cancer. The two plots show the comparison made for the TCGA primary tumours, whereas the two bottom plots show the comparison for the GDSC breast cancer cell lines. The plots on the left column show the upregulated transcripts between each comparison, and that on the right show the downregulated transcripts between each comparison.

Therefore, rather than focusing on the PAM50 classifications of breast cancer cell lines when comparing responses to CSR-targeted drugs, we instead focused on the expression levels of particular drug-targeted CSRs in the cell lines and, for each drug-response test, simply divided the cell lines into higher and lower CSR expression categories (see methods sections). Remarkably, we found that the drug-response profiles of 42% (8 of 19) of the anticancer drugs differed significantly between the higher and lower targeted CSR expression groups (Figure 5, Supplementary File 4). Also, among these drug-response comparisons, 15/19 of the anticancer drugs displayed negative t-values (propensity towards higher efficacy in the higher group than in the lower group), which further confirmed that, for most of these anticancer drugs, drug efficacy is associated with the transcriptional levels of the CSRs that they target.

**Figure 5:**
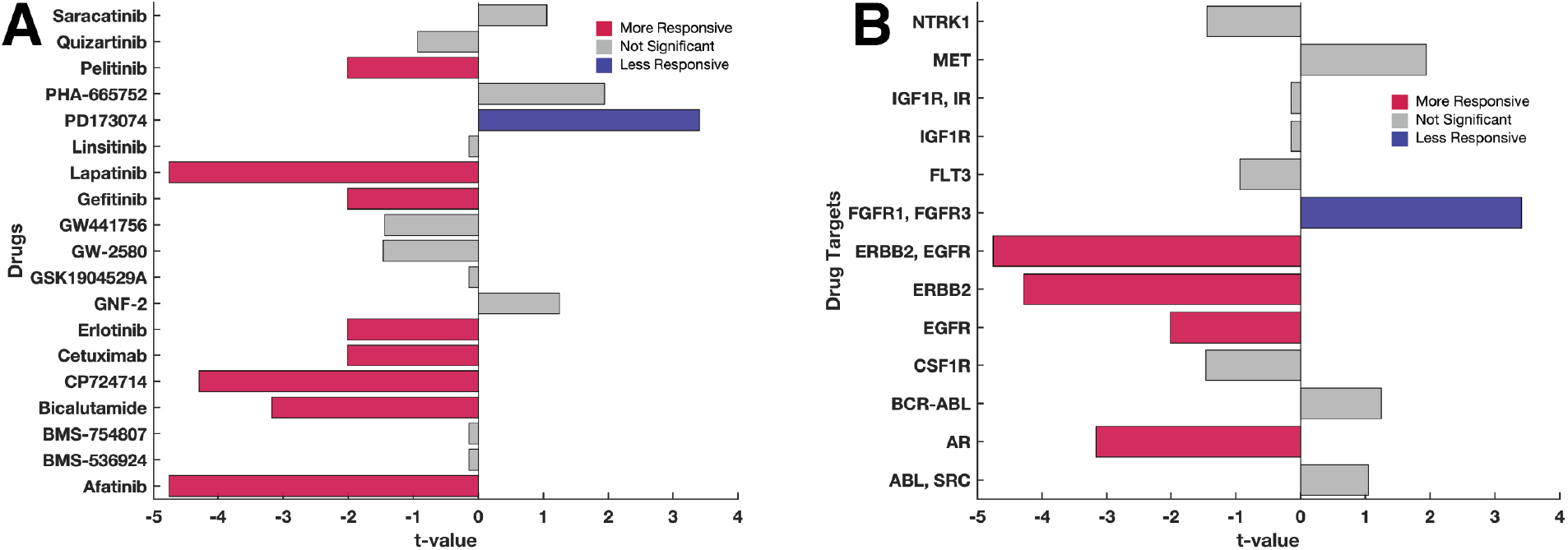
Comparison of the drug-response profiles to CSR-targeting anticancer drugs between the cell lines expressing higher transcript levels of the drug targets and those with lower target transcript levels. Each bar indicates the t-value calculated using the Welch test. The bars are coloured based on the level of statistical significance. Red bars denote statistically significant (p-value < 0.05) increased response to the drug for the cell lines overexpressing the targets compared to those under-expressing the drug target. Grey bars denote no statistically significant difference in the drug response. Blue bars denote statistically significant lowered response to the drug. (A) The comparisons were made across each drug represented in the GDSC database. (B) the comparisons are made across drugs that are grouped based on their target CSRs. For the test statistics relating to each comparison, refer to supplementary file 4.

### The mRNA transcript levels of CSRs in healthy tissues are associated with adverse drug events

We hypothesised that the toxicity of CSR-targeting drugs is associated with the CSR gene transcriptional levels in healthy tissues. Therefore, we examined 224 breast cancer clinical drug trial data by extracting information relating to the anticancer drug tested and reported adverse events that could be ascribed to drug toxicity in specific body tissues for which CSR transcription data was available (see the methods sections and Supplementary File 5).

Next, we segregated the 224 clinical trials into two categories: (1) those involving drugs that target CSRs that are more highly expressed in breast tumours than in healthy body tissues, referred to as “ideal targets” (49 [22%] clinical trials involving 10191 participants); and (2) those involving drugs that target any other CSRs, referred to as the “other targets” (175 [78%] clinical trials involving 40,949 participants). We then compared the proportions of individuals that experienced adverse events between these two clinical trial categories.

We found that the “ideal targets” trials reported significantly fewer drug-associated adverse events than the “other targets” trials (Chi-square test, χ^2^ = 15.2, p-value 9.8 × 10^−5^; Figure 6). Across various categories of adverse effects, a significantly higher proportion of patients in the “other targets” trials (median 4.8%) reported adverse effects than those in the idea targets trials (median = 1.9%; rank-sum test statistic = 186.5; p-value = 0.0094) (Figure 6). Furthermore, we found that the 49 “ideal target” clinical trials utilised 12 unique anticancer drugs, including lapatinib (targeting ERBB2), afatinib (ERBB2 and ERBB4), trastuzumab (ERBB2), and cabozantinib (RET). Conversely, we found that the 175 “other target” clinical trials utilised 74 unique anticancer drugs, among others, filgrastim (targeting CSF3R), etelcalcetide (CASR), and bendamustine (CD69; see Supplementary Figure 2).

**Figure 6:**
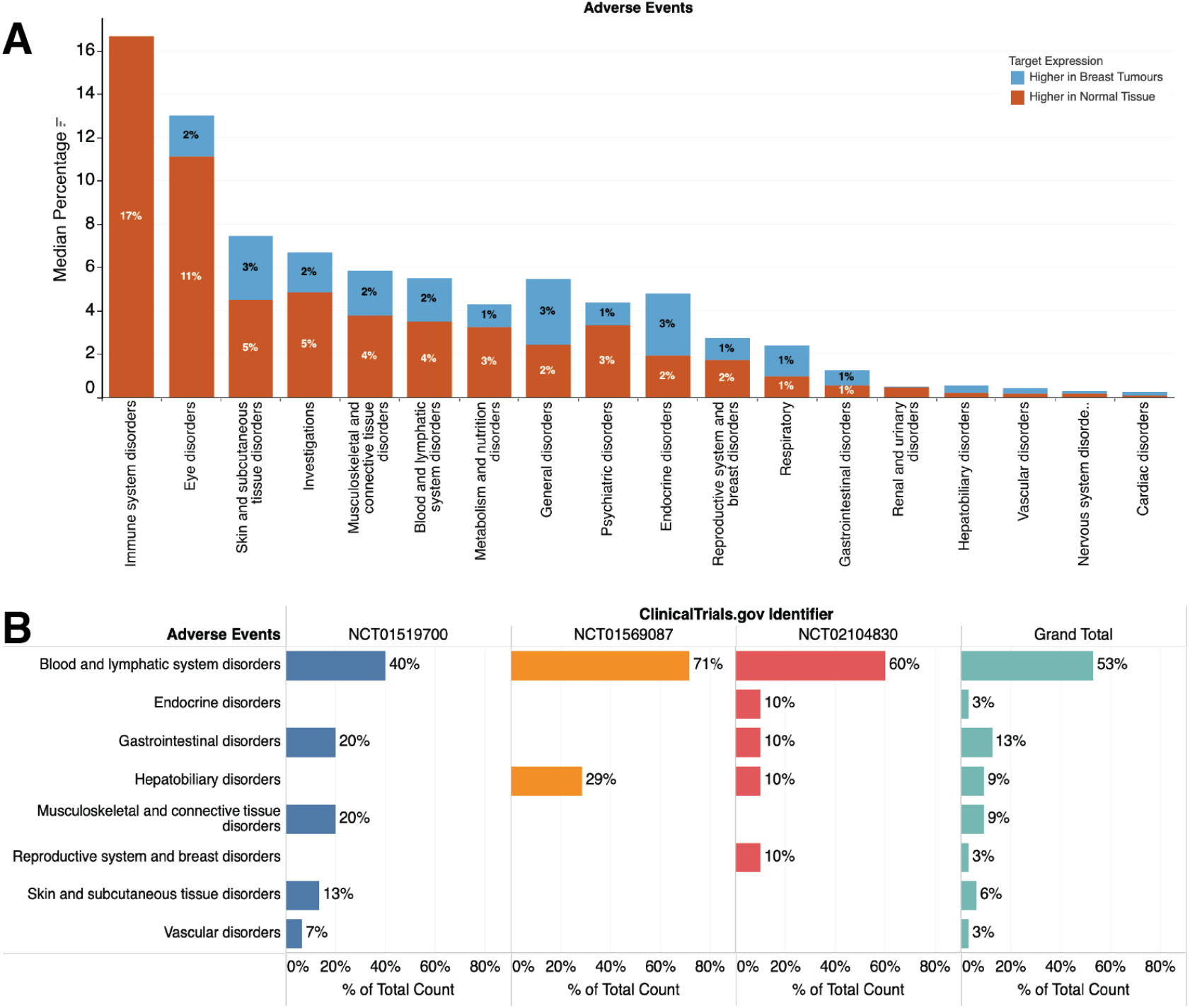
(A) Distribution of adverse events reported in the clinical trial of breast cancer for drugs that target particular CSRs. The group are segregated by the types of drugs that are used to treat breast cancer patients: (1) Those that target highly expressed CSRs (the ideal targets) in breast tumours and (2) those that do not. The median proportion of individuals who experienced adverse events is reported for each bar graph. The colour shows details about adverse events. (B) Percentage of the total count of each category for each reported adverse in clinical trials that utilised the anticancer drug filgrastim, which targets CSF3R.

The anticancer drugs that target the other targets are likely to produce significantly more adverse effects owing to their higher expression in other body tissues compared to the tumour tissues. For example, among the other targets, we found that CSF3R is 338 times more highly expressed in blood cells than in breast tumours (adjusted p-value < 1 × 10^−350^). We analysed the results of three clinical trials (NCT01519700, NCT01569087, and NCT02104830) that utilised the anticancer drug filgrastim, which targets CSR3R and found a high proportion of participants exhibited blood and lymphatic system disorders, which accounted for 17 (53%) of the 32 different reported adverse events (Figure 6B). In clinical trial NCT02104830, anaemia was reported in 44% of the participants, leukopenia (in 95%), and neutropenia (in 95%; Supplementary Figure 3).

Our findings suggest that a positive association may exist between adverse CSR-targeted drug responses and the expression levels of the targeted CSRs in healthy cells.

### Machine learning predicts adverse drug events using the CSR transcription profiles of drug targets across body tissues

We used machine learning methods to determine whether the data from clinical trials could be used to predict the occurrence of adverse drug toxicity events in healthy tissues. Here, we extracted information on tissue-level mRNA transcript measurements for CSRs targeted in published clinical drug trials that reported adverse events affecting healthy tissues (see methods section). We then used these data to train a Gaussian process regression [23] and support vector machine [24] ensemble machine learning model and used this trained model to predict adverse drug effects using an independent test set (see methods section).

For each anticancer drug using tissue-level transcript abundances for the targeted CSR as inputs, our model accurately predicted the healthy tissues that would experience adverse events (R^2^ = 0.75; Figure 7A). Furthermore, for each anticancer drug, the model accurately predicted the proportion of individuals likely to experience adverse events associated with a particular tissue (Figure 7B and 7C; see methods section). For example, in Figure 7B, we show that using the ensemble machine learning model trained on the gene transcript levels of CSRs that are targeted by the drug dasatinib (ABL, SRC, EPH, PDGFR, and KIT), the model predicted the proportion of patients exhibiting an adverse drug reaction (Figure 7), also see Supplementary Figure 4 which show the adverse event prediction for the drug gemcitabine.

**Figure 7:**
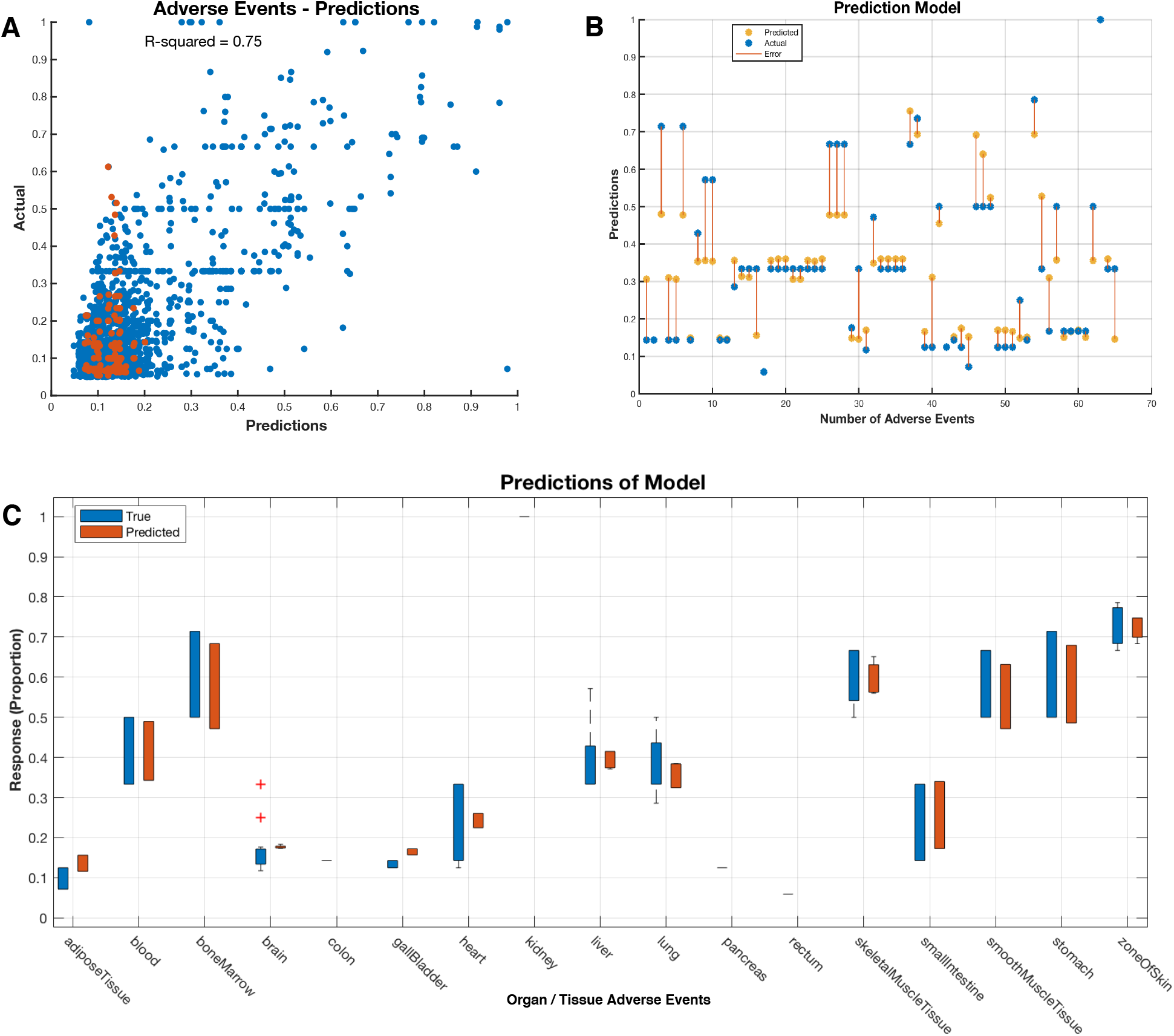
(A) A scatter plot showing the predicted response (y-axis) of the machine learning model plotted against the actual, correct response (x-axis). (B) An example of plots of adverse event prediction for the drug dasatinib. The predicted (blue markers) and actual proportions (orange markers) of individuals that experience adverse events related to a particular organ or body tissue (represented on the x-axis). The line connecting the marker represents the observed error between the predicted proportion of individuals that would experience adverse events against the actual proportion reported in clinical trials. Each prediction is obtained using a model that was trained without using the corresponding (held out) observations reported in breast cancer clinical trials that treated patients with dasatinib. (C) Box plot displaying the typical values of the reported adverse event affecting a particular tissue for the drug dasatinib and the predicted response, and any possible outliers. The central mark indicates the median, and the bottom and top edges of the box are the 25th and 75th percentiles, respectively. The whiskers extend from the boxes to the most extreme data points that are not considered outliers, whereas outliers are shown individually using the “+” symbol.

## Discussion

CSRs are commonly overexpressed in many forms of cancer, making them valuable therapeutic targets for small-molecule inhibitors and antibody therapies [25–28]. Here, we show that the expression of CSR genes varies widely across different healthy body tissues and that this variation can be a significant determinant of adverse anticancer drug reactions. For example, we show that even for CSRs that are more highly expressed in breast cancer tumours than in healthy breast tissue, many of which are currently targeted by breast cancer therapeutics, in some healthy body tissues, many of these CSRs are even more highly expressed than in breast tumours such that these CSR targeting therapeutics could potentially cause considerable collateral damage.

We show that the breast cancer cell lines with higher mRNA transcript levels for CSR genes targeted by a particular drug show a significant tendency to be more sensitive to that drug. While previous studies have also shown that the transcriptional profiles of tumours are correlated with their drug responses [29–32], our focus on CSRs has revealed that to elicit better therapeutic responses and lessen adverse events, “ideal drug targets” would be those CSRs that are both highly expressed in all breast cancer tumours, and poorly expressed in healthy body tissues.

Using published clinical trial data, we showed that treatment regimens targeting CSRs with higher associated transcription levels in breast tumours than any healthy body tissues reported fewer adverse events than those targeting CSRs with associated transcription levels higher in some healthy tissues compared to breast tumours. Recently, another study reported a correlation between the organ systems affected by genetic variations in drug targets and the organ systems in which side effects are observed [15]. This apparent association between the relative transcription levels of CSR genes in tumours and healthy body tissues and adverse effects further emphasises the need to target CSRs that are not only more highly expressed in breast tumour cells than in healthy breast tissues, but which are also expressed at lower levels in other healthy tissues throughout the body.

Therefore, our findings directly apply to selecting CSR drug targets that ultimately yield safer therapeutics. A similar approach may also apply to the identification of non-CSR drug targets that will have minimal toxicity to healthy tissues. Such drug targets would both allow the use of higher drug doses and increase the safety of antibodies conjugated with non-specific cytotoxic chemicals to treat disease [25,33]. The latter is a promising approach that is likely to improve treatment outcomes, as the strategy would not depend on the biological importance of the targeted CSRs within downstream signalling pathways: irrespective of its biological function, any CSR that is expressed in tumour cells but not in healthy cells would be a viable target. This approach to anticancer therapy promises fewer disease remissions and better efficacy than current treatments as non-specifically cytotoxic compounds would not, like current chemotherapeutics, only kill tumour cells with a specific molecular phenotype [34–38]. For as long as the cells in the tumour express the targeted CSR, the non-specific toxin(s) that is(are) delivered to these cells will kill all the cells in their proximity, including those that do not express the CSR.

Altogether, we have identified differentially expressed CSR transcripts between breast tumours and all other healthy body tissues that could maximise the therapeutic benefits while reducing off-target toxic effects of CSR targeting small molecules and immunotherapeutics. We further propose that publicly available data with sufficient detail and depth is now available to adopt an approach such as we have explored here for the selection of non-CSR drug targets too.

## Method

We accessed transcriptome profiles of healthy tissues from the Genotype-Tissue Expression consortium (https://gtexportal.org/home/); 54 body tissue [16,17]. By collating these data, we obtained the 101 unique transcriptome profiles of all major organs and various tissue types in the human body (Supplementary Files 1). Then we curated a list of CSRs using information from the literature, UniProt Knowledgebase [39], Surfaceome database [40], and the Gene Ontology Consortium [41] using the Gene Ontology term plasma membrane (Supplementary File 1). Finally, we used this list of CSRs to extract only the mRNA transcription data of genes that are CSRs, to which we applied unsupervised hierarchical clustering to reveal the clustering of healthy tissues based on their CSR expression (see Figure 8).

**Figure 8:**
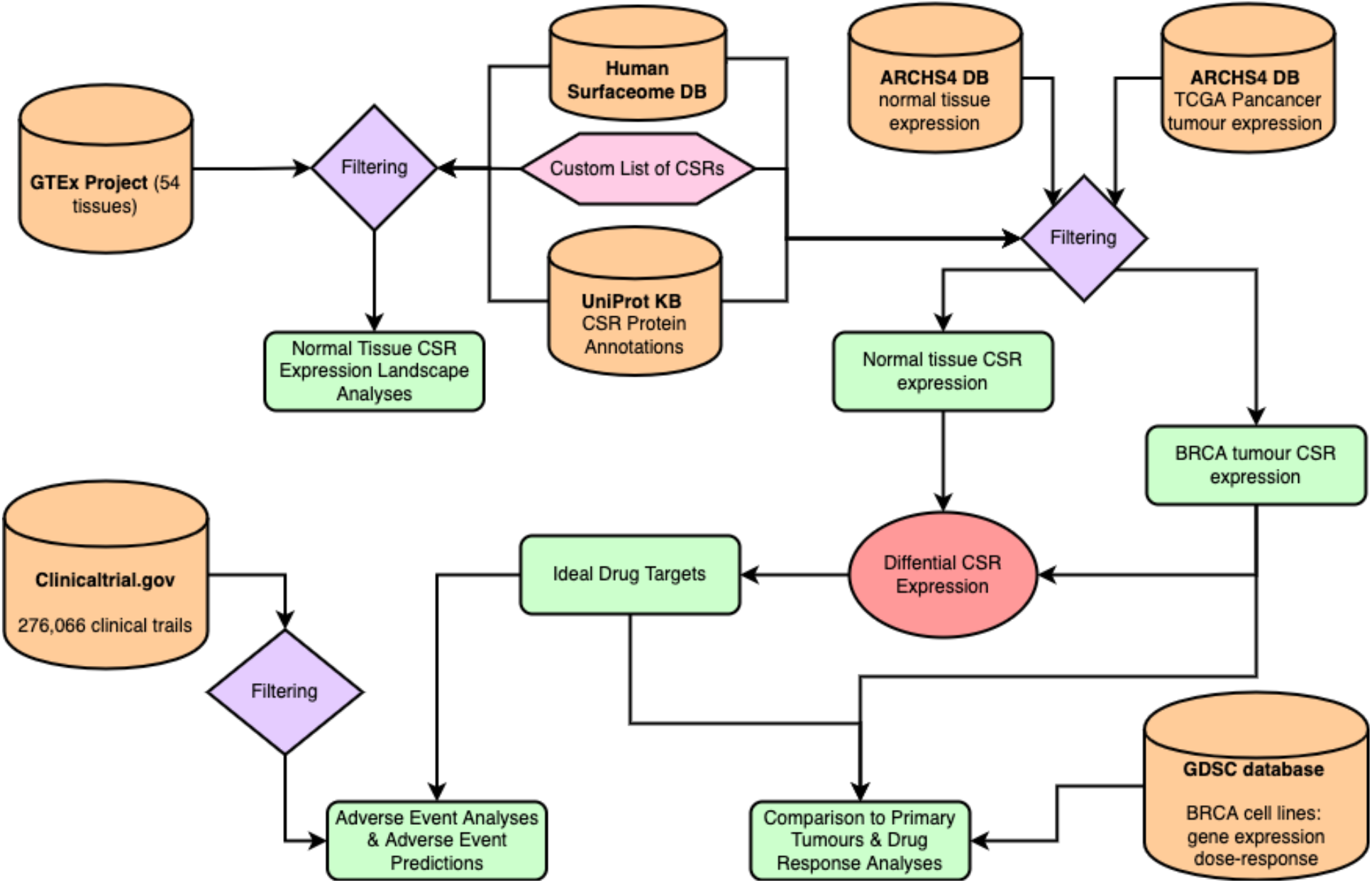
Graphical representation of the overall study method and analysis procedure.

### mRNA expression of CSR across breast tumours and healthy tissues

To establish the landscape expression of CSR across healthy tissues and breast cancer tissues. First, we obtained evenly processed mRNA expression data from 9,685 individuals across 31 healthy tissues compiled by the GTEx and 1,079 breast cancer samples by the TCGA from the ARCHS^4^ database [42,43]. We used these datasets, which are directly comparable because they were re-processed using the same computation pipeline by the reCount2 project [44]. Then, we applied t-Distributed Stochastic Neighbourhood Embedding to reveal the clustering pattern of breast cancer tumours and healthy tissues [45].

### Identification of the ideal drug target CSR

We hypothesised that anticancer drugs that target CSR are also likely to exhibit off-target adverse effects related to CSR expression in other healthy tissues. Therefore, the best drug target CSRs would be those that are upregulated in the disease state compared to any other healthy body tissues. To identify such CSR targets, we performed differential gene expression using the negative binomial test [46], by comparing the CSR transcript between the mRNA transcription levels in breast tumours from TCGA and each healthy tissue from the GTEx project see Supplementary File 2). Then, we return the CSR transcripts that we found upregulated (adjusted p-value < 0.05 and fold change > 2) across all the breast cancer tissues verse healthy tissues comparisons (i.e., the intersection) to yield what we propose as the “ideal” drug and antibody targets (Figure 2).

### Prediction of breast cancer subtypes using their CSR transcription data

Since breast tumours are subtyped using the PAM50 classification (Luminal A, Luminal B, Normal-like, basal and HER-2 positive) scheme, we set out to find out if we reproduce the tumour subtype using only mRNA transcripts of the CSRs. First, using the breast cancer PAM50 subtypes reported by the TCGA for each sample and their transcript levels of CSR genes, we applied an embedded feature selection method based on the boosted decision tree-based machine learning algorithm to identify 50 CSR features that were the most important predictors of the breast cancer PAM50 subtypes. Then we trained an ensemble prediction model by aggregating 20 decision trees [47] using Random Undersampling Boosting [48], which we used to predict the breast cancer subtypes of the TCGA breast tumours into the Luminal A, Luminal B, Normal-like, basal and HER-2 positive subtypes.

### Influence of CSR transcription on the PAM50 subtypes’ response to drugs

To evaluate how the mRNA transcriptions of CSR may be related to the overall response of PAM50 subtypes of breast tumours to an anticancer drug, we utilised the corresponding PAM50 subtype of breast cancer cell lines. Here, we used the dose responses from the GDSC [22] of breast cancer cell lines that were previously classified into the four breast cancer subtypes by Dai et al. [49] and Aniruddha [50]. For each group of cell lines that belong to two different breast cancer subtypes, we compared their drug response to a particular anticancer drug for all 32 anticancer drugs using the student t-test with unequal variance assumed (see Supplementary File 4).

### Comparison of the CSR transcription in primary tumours versus cancer cell lines

To evaluate the mRNA transcription of CSRs of the primary breast cancer PAM50 subtypes represented in the TCGA database with their corresponding PAM50 cancer cell lines that are represented within the GDSC database, we compared the list of the differentially expressed transcript between each pair of PAM50 subtypes. More specifically, first, we identified the differentially expressed CSR transcripts between each pair of PAM50 subtypes of the GDSC cancer cell lines using the Welch test.

Then we compared the list of differentially expressed CSRs between each pair of breast cancer PAM50 subtypes (e.g., basal vs HER2 positive) of the primary tumours and those of the corresponding cancer cell lines (basal vs HER positive; see Supplementary File 3). Here, we expect a concordant list of up and down-regulated for each matched comparison.

### Association of cell line CSR transcription and drug response

We classed the breast cancer cell lines into two categories for each drug-response comparison regardless of their breast tumour PAM50 subtype classification: 1) those that overexpress the CSR target of the drug and 2) those that underexpress the target CSR for the drug. To dichotomise the cancer cell lines into these two groups, first, for each drug, we retrieved the transcription profile of the targets across cell lines from the GDSC and applied z-normalization to the profile [51]. Then using a cut-off value of 1 standard deviation, we categorised the cell lines with z-score values above 1 as those with higher drug target expression and the cell lines with a z-score less than −1 as those with lower drug target expression. The cell lines with z-scores that fall from −1 to 1 were excluded from the comparison on a drug-to-drug basis. To compare the mean drug responses of these two groups of the cell lines (high drug target expression and lower drug target expression groups), we applied a Welch test on the cell line’s group area-under-the-curve of the dose-response curve (see Supplementary File 4)

### Validation of the ideal targets using reported adverse events

We retrieve data from https://clinicaltrials.gov/ of breast cancer clinical trials that applied a single anticancer drug or antibody [52]. Further, we obtained the actual targets of each of the drugs applied in the clinical trials from the Pharos database (https://pharos.nih.gov [53]) and the Drug Gene Interaction Database (http://dgidb.org [54]) to return only clinical trials that utilised drugs that target CSRs. The clinical trial records include information on participants of clinical trials, such as the treatment and adverse events that were experienced by each participant (see Supplementary File 5). Therefore, we extracted information on the anticancer treatments used and the adverse events reported for each anticancer drug.

Then we segregated the clinical trials into two sets: those that utilised CSRs that we identified as the “ideal” targets (i.e., highly expressed in breast cancer compared to any other healthy body tissues) and those that utilised the “other targets”. The clinical trials that employed the “ideal targets” reported adverse events for 544 individual participants, whereas those that employed “other targets” reported adverse for 501 individual participants. Finally, we compared the reported proportions and the actual number of individuals that experienced adverse events between these two categories of clinical trials.

### Prediction of the adverse events using the CSR transcription levels of the healthy tissues

We mapped the adverse events reported in the clinical trials to particular body tissue, e.g., “Skin and subcutaneous tissue disorders” were ascribed to the skin, whereas “cardiac disorders” were ascribed to the heart (see Supplementary File 5). Then we annotated each of these adverse events to the healthy tissue expression of the CSR transcript levels that we obtained from the GTEx data, e.g., the row in the data that specifies the adverse events that occur in the heart due to some anticancer drug are related to the CSR expression of the healthy heart (Supplementary File 5).

Further, for each drug used in the clinical trial, we also obtained the drug target from the Pharos database and the Drug Gene Interaction database to return only clinical trials that utilised drugs that target CSRs [53,54].

We used these data – adverse events ascribed to a particular tissue and the tissue CSR transcript levels of the drug target (use in the treatment and producing the adverse events) – to train a machine learning model that would be used to predict the adverse events in various healthy tissue. Here we trained 20 different machine learning regression models, including linear regression (using a simple linear model, interaction terms and the stepwise methods), decision trees regression (of various tree and leaf sizes), support vector machines regression (of various kernel scale, kernel function and box constraint), ensemble trees (boosted and bagged trees), and Gaussian process regression (of various kernel scale, kernel function and signal standard deviation and sigma). We selected the two best-performing regression models based on the 5-fold cross-validation accuracy: the quadratic support vector machines model [24] (root mean squared error = 0.042) and the squared exponential Gaussian process regression [23] (root mean squared error = 0.043). We have then combined these two best-performing models by training an ensemble machine learning algorithm based on quadratic support vector machine regression and squared exponential Gaussian process regression. We used this ensemble model to predict the adverse events that each particular anticancer would produce based on the CSR expression of the anticancer drug targets in various healthy tissues.

### Statistical analysis and data visualisation

All statistical analyses were performed in MATLAB 2019b. Fisher’s exact test was used to assess associations between categorical variables. In addition, the independent sample Student t-test, Welch test and the one-way Analysis of Variance were used to compare continuous variables where appropriate. Statistical tests were considered significant at p < 0.05 for single comparisons, whereas the p-values of multiple comparisons were adjusted using the Benjamini-Hochberg method [55]. All the results and data were visualised in MATLAB 20109b or Tableau version 2019.1.7.

## Data Availability

The data that support the findings of this study are available from the following repositories: The GTEx portal (https://gtexportal.org/home/), ARCHS^4^ (https://amp.pharm.mssm.edu/archs4/), clinicaltrial.gov (https://clinicaltrials.gov/), Genomics of Drug Sensitivity in Cancer (https://www.cancerrxgene.org/).

## Acknowledgements

Student bursary funding for this project was provided by H3ABioNet, supported by the National Institutes of Health Common Fund under grant number U24HG006941. The content of this publication is solely the responsibility of the authors and does not necessarily represent the official views of the National Institutes of Health. This work is based on the research supported in part by the National Research Foundation of South Africa (Grant Number 47904).

## Ethics Approval

The University of Cape Town; Health Sciences Research Ethics Committee IRB00001938 approved the protocol of this study. This study involved analysing publicly available datasets collected by the TCGA, ICGC, GDSC and other databases from consenting participants. All methods were performed following the relevant policies, regulations and guidelines provided by the TCGA, ICGC, GDSC and other databases for analysing their datasets and reporting the findings.

## Author Contributions

Conceptualisation, SB, MS, and DPM; Methodology, MS, KN, DR, NM, SB, and DPM; Formal Analysis, MS, KN, DR, NM, and DPM; Writing – Original Draft, MS, KN, and DPM; Writing – Review & Editing; MS, KN, DPM, DR, NM, and SB; Visualisation, MS; Supervision, SB, and DPM.

