## Supplemental Information for "A Machine Learning and Bioinformatic Analysis Reveals an Associated between Cell Surface Receptor Transcript Levels with Drug Response of Breast Cancer Cells and the Drug Off-Target Effects"

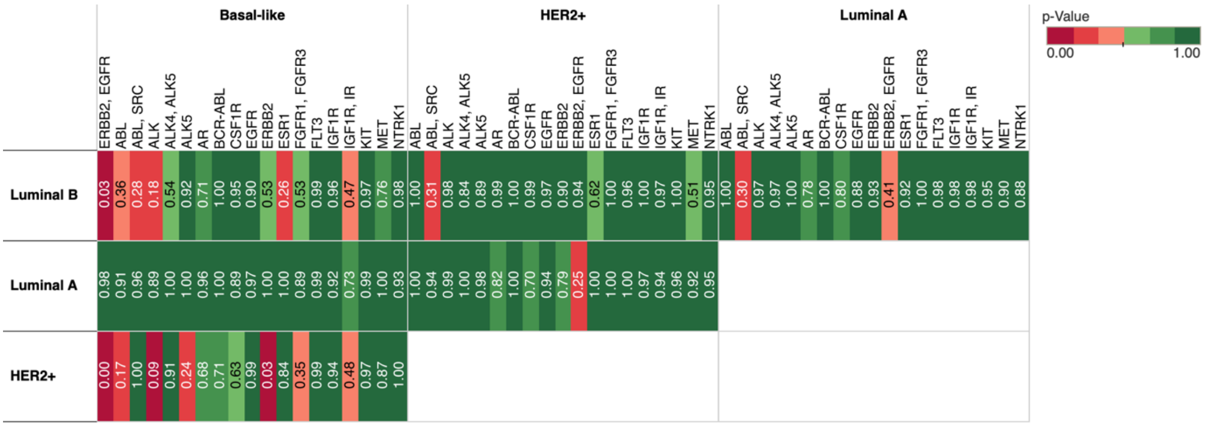


**Figure 1:** Distribution of dose-response comparisons between PAM50 subtypes of breast cancer cell lines. The colours show the degree of statistical significance (i.e., p values), with redder colours denoting smaller (more significant) p-values and greener colours denoting larger p-values. The marks are labelled with p-values.

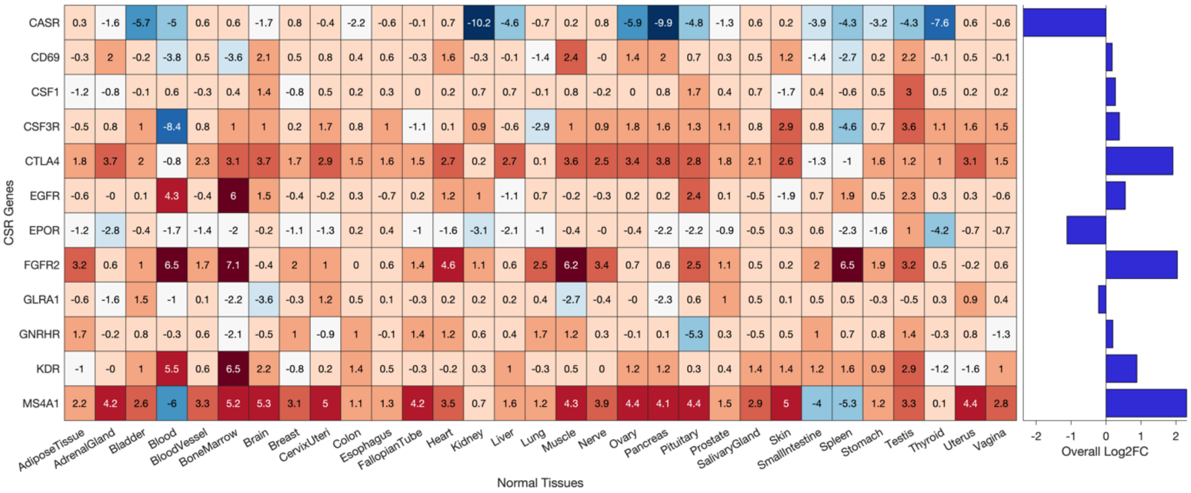


**Figure 2:** The heatmap of the log-2 fold-change values of the CSR transcripts of some anticancer drug targets utilised in the clinical trial data analysed.

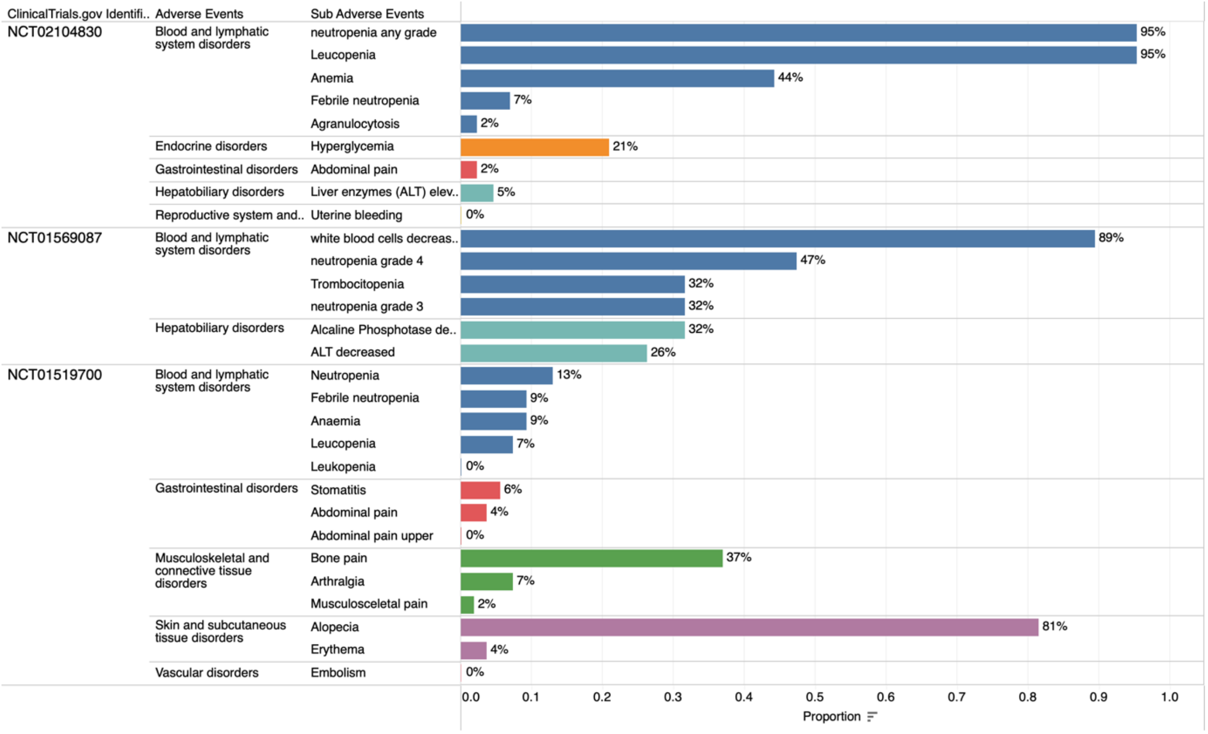


**Figure 3:** Adverse events reported in three clinical trials that utilised the drug filgrastim, which targets CSR3R. Each bar shows the percentage of individuals that experienced the adverse events broken down by sub-adverse events for each clinical trial denoted by ClinicalTrials.gov. The colours show details about adverse events categories.

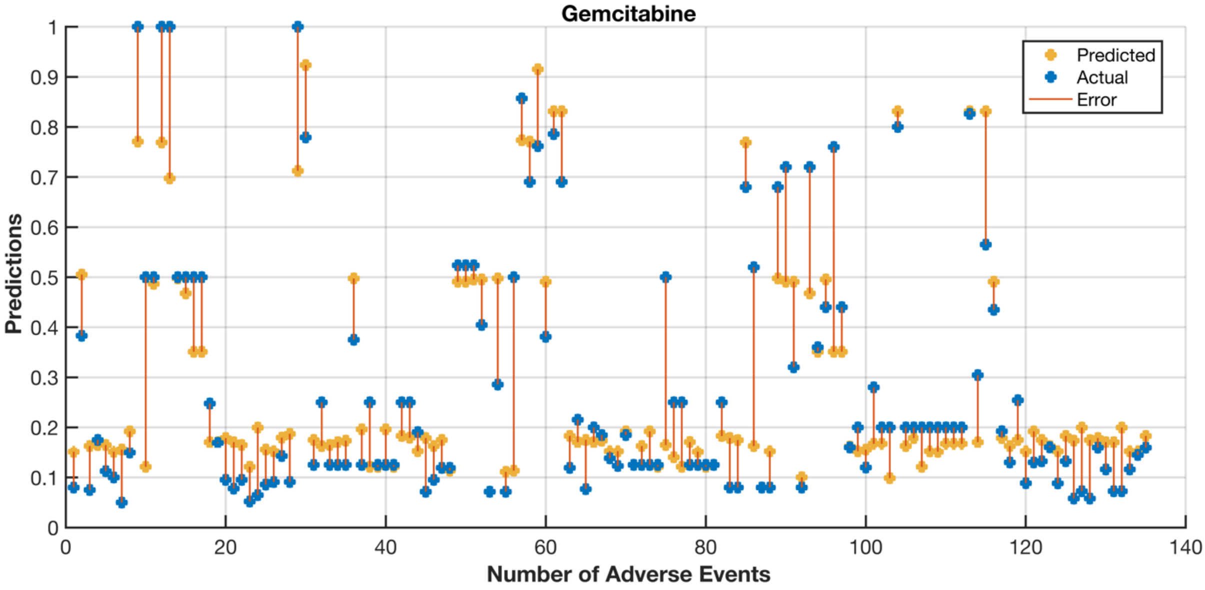


**Figure 4:** Adverse event prediction for the drug gemcitabine. The predicted (blue markers) and actual proportions (orange markers) of individuals that experience adverse events related to a particular organ or body tissue (represented on the x-axis). The line connecting the marker represents the observed error between the predicted proportion of individuals that would experience adverse events against the actual proportion reported in clinical trials. Each prediction is obtained using a trained model without using the corresponding (held out) observations reported in breast cancer clinical trials that treated patients with gemcitabine.

**Supplementary File 1:** Data of differentially expressed CSRs between breast cancer and all other healthy body tissues and organs: The spreadsheet contains the following results according to the sheet name. ***Up Genes - BRCA vs All Normal***; list of genes we found highly expressed in breast cancer than in any other healthy body tissue. ***Up Genes Counts- BRCA vs Normal***; the number of upregulated genes between breast cancer and each healthy tissue, and the number of downregulated genes between breast cancer and each healthy tissue.

**Supplementary File 2:** All differentially expressed CSRs between breast cancer and each healthy tissue. Each sheet is named according to the comparison for which the differential CSR expression results are represented. E.g., BRCA vs skin shows the results between breast tumour profile by the TCGA[1] and all healthy tissues profile by the GTEx project[2].

**Supplementary File 3:** Differentially expressed CSRs between PAM50 subtypes of breast cancer. The sheets are named according to the comparison for which the differential expression CSRs results are represented. E.g., BRCA Basal-Vs-BRCA HER2 shows the results of Basal-like breast cancer subtype and HER2-positive breast cancer.

**Supplementary File 4:** Drug-Response Differences. The spreadsheet contains the following results according to the sheet name. ***Dose-Response of cell lines***; collated data of breast cancer cell lines profiled by the GDSC [3,4] and the PAM50 subtype of each breast cancer cell line. ***Dose Response Anova Results***; comparison of dose-response between each PAM50 subtype of breast cancer. ***Anova Statistics***; ANOVA statistic for the comparison in the "Dose Response Anova Results" spreadsheet. ***CSR_based Response Comparison***; Dose-response comparison between the GDSC breast cancer cell line that we segregated into two groups, those that expressed higher amounts of a particular drug target and those that expressed lower amounts of a particular drug target.

**Supplementary File 5:** Spreadsheet showing the information that we collated from various results, including clinical trials information from www.clinicaltrials.gov of drug targets for drugs used applied in clinical trials that were retrieved from the Pharos database and Drug Gene Interaction database and the CSR transcript levels of the tissues in which are adverse events that are reported in the clinical trials occurred.
